# LeukGenePipeline: Modular Workflow for Genomic Datasets

**DOI:** 10.1101/2025.08.01.668175

**Authors:** Ana Carolina Pacífico dos Santos, Omar Arias-Gaguancela

## Abstract

Analyzing human genome data has become increasingly common, supported by the growing availability of public repositories that enable statistical modeling and predictive analysis. However, researchers from wet lab-based disciplines often face challenges due to limited training in programming and computational tools. To address this barrier, we introduce LeukGenePipeline (LGP): a user-friendly, Python-based tool designed to automate core genomic analyses. As a proof of concept, LGP was used to perform mutation classification, copy number variation (CNV) analysis, pathway enrichment analysis (PEA), and gene ontology (GO) enrichment using data from the COSMIC public database (v101) with a focus on acute myeloid leukemia (AML). Data consisted of a mutation table with 830,978 unique rows associated with protein-coding genes, and a CNV table with 12,926 gene-level entries. LGP outputs revealed frequently mutated, CNV-altered genes, and enrichment of key transcription factors associated with leukemogenesis.

## Introduction

Advances in genomic technologies have created unprecedented opportunities to improve the diagnosis, treatment, and management of genetic disorders [1]. As bioinformatics tools continue to evolve, efficiently managing and analyzing human genome data has become crucial to sustaining research productivity and extracting meaningful insights from high-throughput datasets [2].

One common approach involves the use of public genomic databases, which typically requires a multi-step process: (1) identifying relevant datasets and their sources, (2) obtaining access permissions, (3) downloading the data, (4) annotating and characterizing the content, and (5) evaluating data quality for downstream applications [3].

However, after data acquisition, researchers, especially those without a computational background, often face difficulties in performing integrative and large-scale analyses [4,5]. In the context of leukemia research, integrative analyses that combine mutation profiles [6], copy number variations (CNVs) [7], and functional enrichment [8] are critical molecular mechanisms involved in leukemogenesis [6,7]. Yet, the preprocessing, cleaning, and classification steps can be time-consuming and technically demanding [9]. This creates a barrier for many scientists who may lack the training required to perform such analyses.

To address this gap, we developed LeukGenePipeline (LGP), a user-friendly Python-based tool, designed to automate core genomic analyses in leukemia datasets (Figure 1). Mutation and CNV data were retrieved from the COSMIC (Catalogue Of Somatic Mutations In Cancer) database, one of the largest manually curated repositories of cancer-related somatic alterations [23].

**Figure 1:**
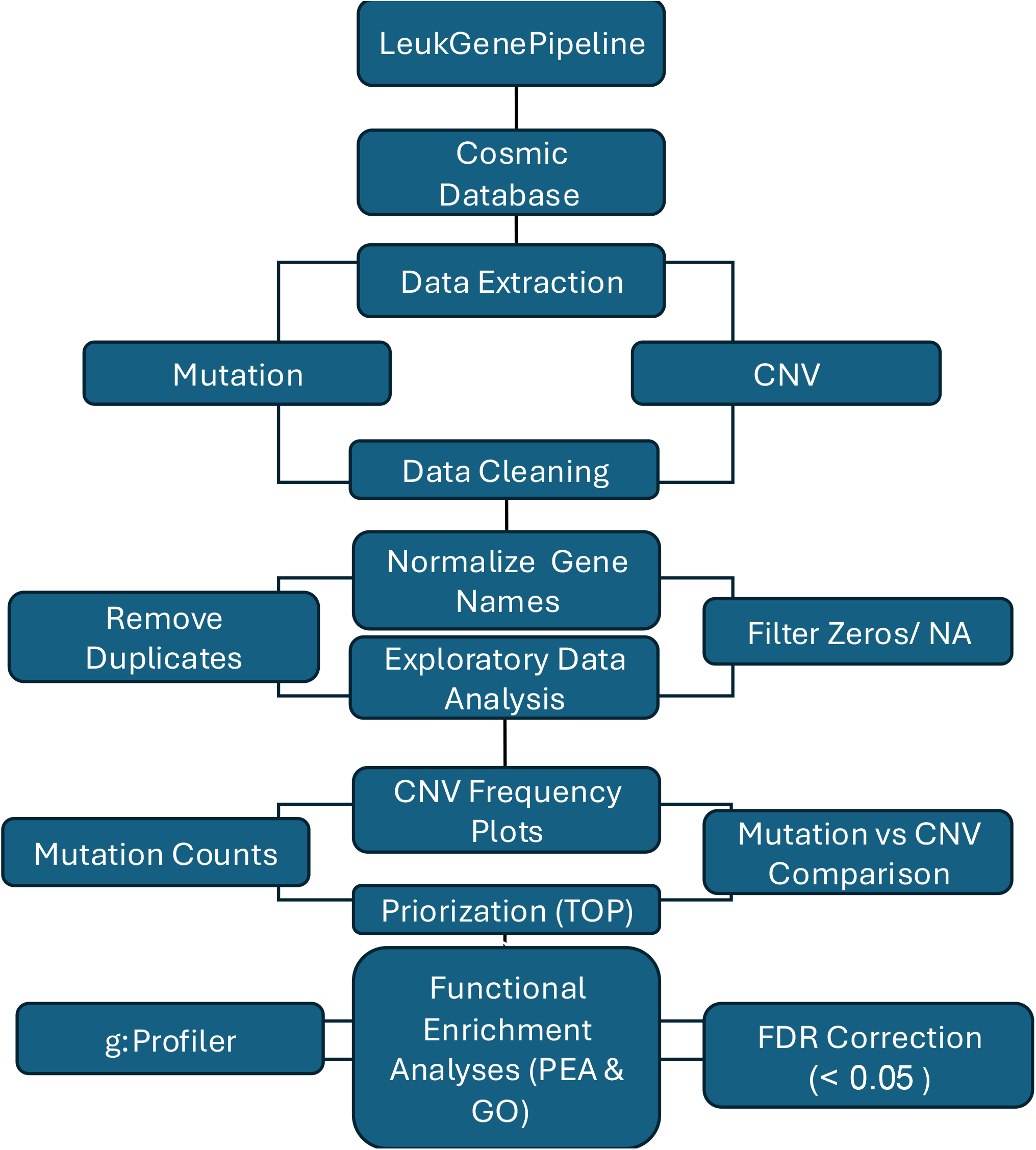
LeukGene Pipeline. After data mining from the COSMIC database, the mutation and CNV datasets were cleaned and standardized through a quality control procedure. Exploratory analyses included gene-wise and alteration-type distributions. Genes with the highest CNV frequency were subsequently used for pathway enrichment analysis (PEA) and gene ontology (GO) annotation.

LGP uses a formatted CSV file as input, and by simply running the notebook, users can perform mutation classification, CNV quantification, Pathway Enrichment Analysis (PEA), and Gene Ontology (GO) enrichment with minimal manual intervention. As an open-source resource, the notebook can also be easily customized or extended by users to fit specific datasets or research goals. All outputs, including structured tables and high-resolution figures, are automatically saved in dedicated folders for downstream interpretation and reporting. By streamlining the analysis workflow and removing technical barriers, LGP supports more standard research practices and helps researchers focus on biological interpretation rather than data handling.

## Method

Mutation and copy number variation (CNV) data were obtained from the Catalogue of Somatic Mutations in Cancer (COSMIC), a curated database of somatic variants across human cancers. The mutation file included columns such as Gene Name, Mutation CDS, and Mutation AA, The CNV dataset contained fields like Gene, Gain, High-level Gain, Total Samples, Loss, High-level Loss, No Change, Net CNV, and Total CNV Event. To aid user understanding, illustrative subsets of the datasets are provided in Supplementary Tables S4 and S5, showing the structure and sample entries from each file. These examples reflect the raw format before any preprocessing. Datasets were downloaded as CSV files and filtered to retain only entries related to hematologic malignancies (e.g., leukemias).

A harmonized gene identifier format was adopted to ensure that both CNV and mutation datasets could be merged accurately for integrative visualization and statistical modeling. Datasets were downloaded as CSV files and filtered to retain only entries related to AML.

Non-coding or ambiguous annotations present in COSMIC exports were excluded by retaining only rows corresponding to protein-coding genes. Code and example outputs are available on a GitHub repository (https://github.com/CarolPacifico0/LeukGenePipeline) to support transparency and reproducibility of the workflow.

All outputs from these notebooks, including tables and figures, will be saved in dedicated folders. The resulting spreadsheet tables will be saved in a folder called ‘TABLES’ while the generated figures will be saved as TIFF images in the folder “PLOTS” (Table S1).

## Preprocessing

During the preprocessing phase, raw genomic data were filtered and reformatted to enable robust downstream analyses. This general structure aligns with standard bioinformatics workflows for variant data preprocessing [10], but the specific steps applied in this pipeline were custom-designed for this project, as described below:

The mutation dataset was initially cleaned by removing entries with missing gene names and non-standardized gene identifiers. To ensure compatibility with downstream annotation and enrichment tools, all gene names were stripped of transcript-specific tags (e.g., ENST Ids) and converted to uppercase standardized gene letters. The initial file contained 831,839 entries. After removing an invalid header row and applying quality control filters (missing values, duplicate rows, and transcript-specific IDs), 830,978 valid entries were retained. Additionally, redundant rows were eliminated, and amino acid alterations were classified into seven categories: missense, synonymous, nonsense, insertion, deletion, frameshift, and unknown. This classification was used in the exploratory stage to understand the distribution of mutation types and select genes for mutation-CNV comparative analysis.

For the CNV dataset, a similar preprocessing pipeline was applied. Rows lacking gene identifiers were excluded, and duplicate entries were removed. For CNV data, no missing values were found in the gene column, and the initial 12,926 entries were preserved after quality control and column standardization. The dataset was then scanned for structural variation types: insertion, inversion, translocation, or duplication, but none were found. As a result, the CNV analysis focused solely on four canonical types of alterations: gain, high-level gain, loss, and high-level loss. Numeric columns were coercively converted to float format to ensure compatibility with aggregation functions, and genes with a total CNV count of zero were filtered out before enrichment analysis.

## Quality Control (QC)

Several quality control checks were implemented to ensure data consistency and biological interpretability. This included detection and removal of missing values in key columns such as gene names, mutation types, and CNV event counts; filtering out rows with zero CNV or mutation counts, as these are not informative for downstream analyses; standardization of gene names by removing whitespace and converting all entries to uppercase to ensure consistency across datasets; elimination of duplicated entries to avoid redundancy. These steps were essential to ensure data harmonization before proceeding with exploratory analysis and pathway-based methods.

In the mutation data, samples were reviewed for high-impact alterations (e.g., frameshift mutations, stop-gain variants) and artifacts, and only entries associated with protein-coding genes were retained.

For the CNV dataset, gene-level CNV burden was calculated by summing the counts of gain, high-level gain, loss, and high-level loss. Genes were ranked by their total CNV burden (sum of gain, high-level gain, loss, and high-level loss). The top 200 genes were selected for enrichment analyses.

Genes were ranked based on their total CNV burden, calculated as the sum of gain, high-level gain, loss, and high-level loss events. The genes with the highest cumulative CNV event counts were selected for enrichment analysis, rather than using an absolute threshold. This approach ensures that only genes with the most pronounced genomic alterations were retained for downstream interpretation.

The integrity of gene identifiers across both datasets was checked to ensure there was no mismatch during the integration stage. Final datasets were re-validated by confirming the absence of missing values in key columns and by generating summary statistics and visual inspections (e.g., CNV type distribution plots and mutation frequency histograms).

## Outputs

We used LeukGenePipeline to perform mutation classification, CNV detection, pathway enrichment (PEA), and gene ontology (GO) enrichment on leukemia genomic data (Figure 1).

In the Mutation Analysis notebook, 830,978 mutations were classified into seven categories: missense, nonsense, synonymous, frameshift, insertion, deletion, and unknown. As illustrated in Figure 2A, excluding the “unknown” category, missense mutations were the most frequent class, consistent with established patterns in leukemia genomes [10]. These variants often disrupt protein structure and stability, contributing to specific molecular phenotypes in acute leukemia [11]. Figure 2B highlights the top 10 most frequently mutated genes, including key leukemia-associated genes such as TP53 [12] and FLT3 [13]. This aligns with known mutational subsets in acute myeloid leukemia (AML) [14].

**Figure 2:**
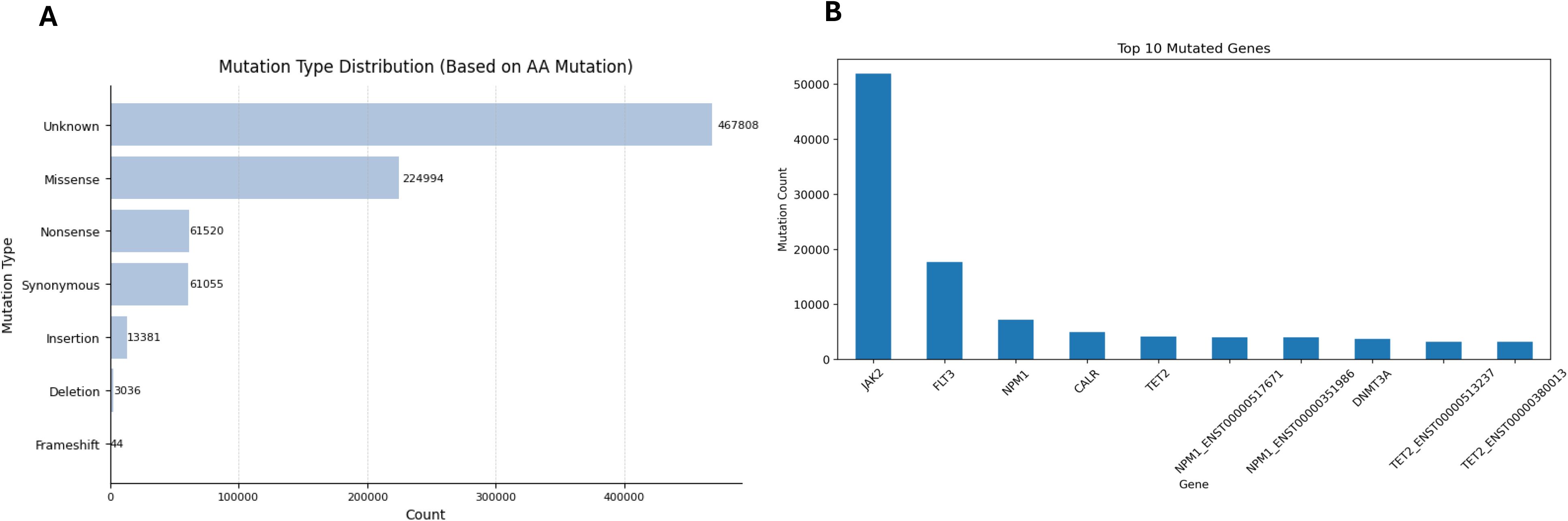
Mutation type classification. Upon preprocessing, mutation events were classified based on amino acid changes. A) Distribution of mutation types: unknown, missense, nonsense, synonymous, insertion, deletion, and frameshift. B) Top ten most frequently mutated genes in the dataset

The second notebook analyzed CNV data from the same cohort. After quality control, a total of 12,926 gene-level entries were retained for analysis. As shown in Figure 3A, the top 10 genes with the highest CNV burden included COX4I2 and DUSP15, both previously implicated in AML pathogenesis [15,16].

**Figure 3:**
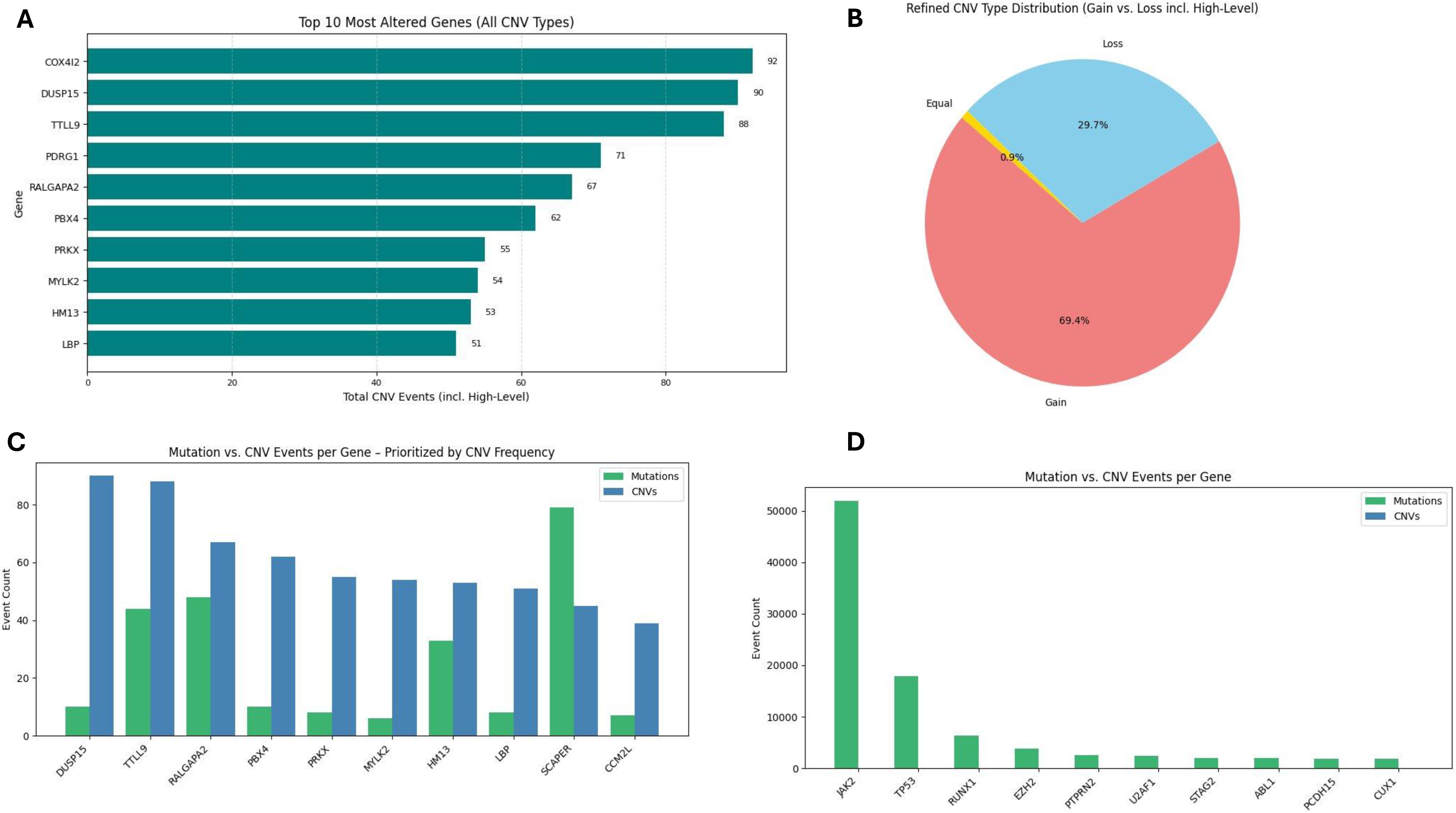
Combined analysis of copy number variation analysis (CNV) and point mutations. A) Top ten genes with the highest CNV frequency, considering gain, high-level gain, loss, and high-level loss. B) Refined CNV-type distribution. C) Side-by-side comparison of mutation and CNV counts per gene. D) Overlap between the most frequently mutated genes and their corresponding CNV alterations, including detailed breakdown by CNV type.

Figure 3B displays the classification of CNV events into four types: gain, high-level gain, loss, and high-level loss. The majority (69.4%) were gains, which are often associated with hyperdiploidy in acute lymphoblastic leukemia (ALL) [17] and chronic lymphocytic leukemia (CLL) [16]. Losses accounted for 29.7% and have been linked to recurrent deletions of specific chromosomes and chromosomal regions in AML [17]. Only a small proportion of genes exhibited balanced CNVs (equal gains and losses).

Figures 3C-D offer an integrative comparison of mutation and CNV frequencies per gene. This highlights genes like DUSP15 and TTLL9 as potentially co-altered targets, both of which have been linked to epigenetic changes such as hypermethylation in acute lymphoblastic leukemia (ALL) [15].

The top CNV-altered genes (Table S1) were then used for pathway enrichment analysis (PEA) and GO enrichment (Figure 4; Table S2 and S3). All enrichment results were adjusted using False Discovery Rate (FDR) correction with a threshold of *p* < 0.05. Figure 4A shows the top 10 enriched pathways, featuring transcription factors such as HES-1, ELK1, and SP2, which have been reported as relevant markers in CML and AML [18–20]. Figure 4B presents the top enriched GO terms, including leukemogenesis regulators like HES-1, ER71, and E2F7-well-known for their roles in cell cycle regulation, hematopoietic differentiation, and cellular proliferation[17,20–22]. These findings support the hypothesis that CNVs frequently disrupt transcriptional regulation in leukemia and may help pinpoint functionally relevant genomic regions for further investigation.

**Figure 4:**
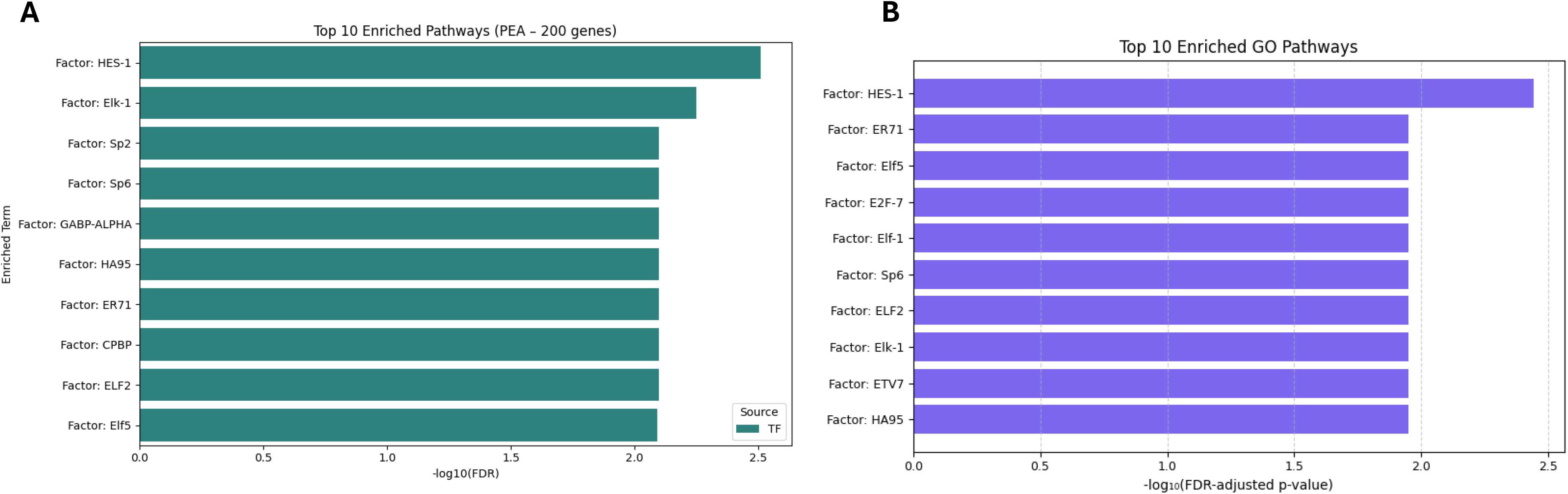
Pathway Enrichment Analysis (PEA) and Gene Ontology (GO). A) Top ten enriched pathways using the 200 most CNV-altered genes. B) Top ten GO terms significantly enriched in the same gene set. They were adjusted using the Benjamini-Hochberg method with a significance threshold set at FDR < 0.05. For access to the complete list of genes used in the enrichment analysis, refer to Supplementary Table S2 and S3.

## Conclusion

In this work, we presented LeukGenePipeline (LGP), a modular and user-friendly computational workflow developed to support exploratory and functional genomic analyses in leukemia. By integrating mutation classification, CNV burden analysis, and pathway and gene ontology enrichment, the pipeline enables rapid interpretation of publicly available datasets with minimal programming effort. The pipeline was tested with curated data from the COSMIC database and enabled the identification of frequently mutated and CNV-altered genes, enrichment of key transcription factors, and gene ontology terms related to leukemogenesis. Overall, we introduced a framework for genome-wide analyses that can be extended or adapted to other cancer types or genomic datasets.

## Acknowledgments

The authors did not receive support from any organization for the submitted work.

## Data Availability Statement

The dataset was obtained through COSMIC’s Mutation, version v101, and CNV export utility without association to a single accession ID. Data were filtered to include only entries related to hematologic malignancies (e.g., leukemia).

Notebooks and all supplementary tables were deposited in the GitHub repository https://github.com/CarolPacifico0/LeukGenePipeline.

LGP used the directory change and mounting to the MyDrive module from QuickProt Notebooks [https://github.com/OmarArias-Gaguancela/QuickProt].

## Supplementary Material

### Supplementary Tables

**Supplementary Table S1**. List of the genes with the highest CNV burden, selected for pathway enrichment analysis (PEA).

**Supplementary Table S2**. Full PEA results derived from the top 200 CNV-altered genes, adjusted using the Benjamini-Hochberg method with a significance threshold set at FDR < 0.05.

**Supplementary Table S3**. Gene Ontology (GO) enrichment analysis results for the same gene set, including FDR-adjusted *p*-values.

**Supplementary Table S4**. Exemplifies the mutation dataset, highlighting fields such as Gene Name, Mutation CDS, and Amino acid change.

**Supplementary Table S5**. Provides an overview of the CNV file, showing gene-level counts for Gene, Gain, High-level Gain, Total Samples, Loss, High-level Loss, No Change, Net CNV, and Total CNV Event.

## References

[1] Howley C, Haas MA, Al Muftah WA, et al. The expanding global genomics landscape: Converging priorities from national genomics programs. Am J Hum Genet. 2025;112(4):751–763. doi:10.1016/j.ajhg.2025.02.008

[2] Tanjo, T., Kawai, Y., Tokunaga, K. et al. Practical guide for managing large-scale human genome data in research. J Hum Genet 66, 39–52 (2021). 10.1038/s10038-020-00862-1

[3] Learned, K., Durbin, A., Currie, R. et al. Barriers to accessing public cancer genomic data. Sci Data 6, 98 (2019). 10.1038/s41597-019-0096-4

[4] Benilton S. Carvalho, Gabriella Rustici, The challenges of delivering bioinformatics training in the analysis of high-throughput data, Briefings in Bioinformatics, Volume 14, Issue 5, September 2013, Pages 538–547, 10.1093/bib/bbt018

[5] Morrison-Smith, S., Boucher, C., Sarcevic, A. et al. Challenges in large-scale bioinformatics projects. Humanit Soc Sci Commun 9, 125 (2022). 10.1057/s41599-022-01141-4

[6] Su Y, Han Z, Ji Y, Liu A, Zou D, Yan L, Liu D, Zhang Z, Wang QF. Patterns and variations of copy number alterations in acute myeloid leukemia: insights from the LeukAtlas database. Leukemia. 2025 Apr;39(4):827–836. doi: 10.1038/s41375-025-02514-9. Epub 2025 Feb 2. PMID: 39894867.

[7] Su Y, Han Z, Ji Y, Liu A, Zou D, Yan L, Liu D, Zhang Z, Wang QF. Patterns and variations of copy number alterations in acute myeloid leukemia: insights from the LeukAtlas database. Leukemia. 2025 Apr;39(4):827–836. doi: 10.1038/s41375-025-02514-9. Epub 2025 Feb 2. PMID: 39894867.

[8] Jian J, Yuan C, Hao H. Identifying key genes and functionally enriched pathways in acute myeloid leukemia by weighted gene co-expression network analysis. J Appl Genet. 2025 May;66(2):347–362. doi: 10.1007/s13353-024-00881-0. Epub 2024 Jul 9. PMID: 38977582.

[9] National Research Council (US) Board on Biology; Pool R, Esnayra J, editors. Bioinformatics: Converting Data to Knowledge: Workshop Summary. Washington (DC): National Academies Press (US); 2000. Maintaining the Integrity of Databases. Available from: https://www.ncbi.nlm.nih.gov/books/NBK44940/

[10] Forbes SA, Beare D, Boutselakis H, Bamford S, Bindal N, Tate J, Cole CG, Ward S, Dawson E, Ponting L, Stefancsik R, Harsha B, Kok CY, Jia M, Jubb H, Sondka Z, Thompson S, De T, Campbell PJ. COSMIC: somatic cancer genetics at high-resolution. Nucleic Acids Res. 2017 Jan 4;45(D1):D777–D783. doi: 10.1093/nar/gkw1121. Epub 2016 Nov 28. PMID: 27899578; PMCID: PMC5210583.

[11] Kikkawa K, Matsuda T, Fujimuro M, Sekine Y. The Atypical Dual Specificity Phosphatase DUSP15 Regulates Jak1-Mediated STAT3 Activation. Biol Pharm Bull. 2024;47(9):1487–1493. doi: 10.1248/bpb.b24-00314. PMID: 39261048.

[12] Shahzad, M., Amin, M.K., Daver, N.G. et al. What have we learned about TP53-mutated acute myeloid leukemia?. Blood Cancer J. 14, 202 (2024). 10.1038/s41408-024-01186-5

[13] Gutierrez-Camino, A., Richer, C., Ouimet, M. et al. Characterisation of FLT3 alterations in childhood acute lymphoblastic leukaemia. Br J Cancer 130, 317–326 (2024). 10.1038/s41416-023-02511-8

[14] Kiyoi H, Kawashima N, Ishikawa Y. FLT3 mutations in acute myeloid leukemia: Therapeutic paradigm beyond inhibitor development. Cancer Sci. 2020;111(2):312–322. doi:10.1111/cas.14274

[15] Navarrete-Meneses MP, Pérez-Vera P. Epigenetic alterations in acute lymphoblastic leukemia. Biomedicine & Pharmacotherapy. 2018;98:886–894. doi:10.1016/j.bmhime.2018.01.004.

[16] Zhang L, Li M, Chan AKN, Liu Q, Kang H, Pokharel SP, Mattson N, Singh P, Yang L, Chen CWD. COX4I1 controls mitochondrial electron transport chain complex IV assembly and leukemia progression in acute myeloid leukemia. Blood. 2023;142(Suppl 1):602. doi:10.1182/blood-2023-188526.

[17] Andersson, J., Aydın, E., Gunnarsson, R. et al. Characterizing the allele-specific gene expression landscape in high hyperdiploid acute lymphoblastic leukemia with BASE. Sci Rep 14, 23181 (2024). 10.1038/s41598-024-73743-8

[18] Tian C, Tang Y, Wang T, et al. HES1 is an independent prognostic factor for acute myeloid leukemia. Onco Targets Ther. 2015;8:899-904. Published 2015 Apr 22. doi:10.2147/OTT.S83511

[19] Guo D, Zhang A, Suo M, Wang P, Liang Y. ELK1-Induced upregulation of long non-coding TNK2-AS1 promotes the progression of acute myeloid leukemia by EZH2-mediated epigenetic silencing of CELF2. Cell Cycle. 2023 Jan;22(1):117–130. doi: 10.1080/15384101.2022.2109898. Epub 2022 Aug 8. PMID: 35941836; PMCID: PMC9769447.

[20] Vivian G. Oehler, Ka Yee Yeung, Yongjae E. Choi, Roger E. Bumgarner, Adrian E. Raftery, Jerald P. Radich; The derivation of diagnostic markers of chronic myeloid leukemia progression from microarray data. Blood 2009; 114 (15): 3292–3298. doi: 10.1182/blood-2009-03-212969

[21] Sumanas S, Gomez G, Zhao Y, Park C, Choi K, Lin S. Interplay among Etsrp/ER71, Scl, and Alk8 signaling controls endothelial and myeloid cell formation. Blood. 2008 May 1;111(9):4500–10. doi: 10.1182/blood-2007-09-110569. Epub 2008 Feb 12. PMID: 18270322; PMCID: PMC2343590.

[22] Salvatori, B., Iosue, I., Mangiavacchi, A. et al. The microRNA-26a target E2F7 sustains cell proliferation and inhibits monocytic differentiation of acute myeloid leukemia cells. Cell Death Dis 3, e413 (2012). 10.1038/cddis.2012.151

[23] Wellcome Sanger Institute. (n.d.), v101. Catalogue of Somatic Mutations in Cancer (COSMIC). Retrieved April 21, 2025, from https://cancer.sanger.ac.uk

